# Copper stress upregulates oxidative stress response, histidine production and iron acquisition genes in *E. coli*

**DOI:** 10.64898/2026.03.12.711415

**Authors:** Hanna Ainelo, Karola Jõearu, Andres Ainelo, Angela Ivask

## Abstract

Copper is widely used as a fast-acting antimicrobial, yet the strategies that allow bacteria to survive copper stress remain incompletely understood. Here, we characterize the transcriptional responses of *Escherichia coli* MG1655 to excess ionic copper using RNA sequencing and a genome-wide GFP-based promoter library. We applied 2 mM copper, which slows growth, and 8 mM copper, a near-lethal concentration. RNA-seq revealed extensive transcriptome remodeling, with 487 genes upregulated at 2 mM and 364 at 8 mM. Both concentrations strongly induced canonical copper-responsive systems, oxidative stress defenses, histidine biosynthesis, and multiple iron acquisition pathways – including enterobactin biosynthesis and transport – despite external iron failing to reduce copper toxicity. At 2 mM copper, additional pathways were activated, including heat-shock and protein-folding functions as well as lipid A, methionine and arginine biosynthesis. Copper exposure also repressed large gene sets: 486 genes at 2 mM, enriched for biofilm formation and pH elevation, and 217 genes at 8 mM, enriched for anaerobic metabolism. In contrast to the robust RNA seq results, we investigated the Horizon Discovery *E. coli* genome-wide GFP based promoter library as an alternative screening tool. However, in our experiments it showed low signal to noise ratios, limiting its suitability for large scale gene expression screening.

## Introduction

Copper (Cu) is an essential transition metal required for correct folding and function of a variety of proteins in bacteria, yet it becomes highly toxic when present in excess. Even the mammalian immune system exploits copper’s toxicity by mobilizing it to kill invading bacteria (Focarelli *et al*., 2022). The antibacterial activity of copper has been used for centuries by humans and even now, copper is considered the most effective compound to prevent healthcare-associated infections (Salgado *et al*., 2013; Sifri *et al*., 2016; Zerbib *et al*., 2020; Aillon-Garcia *et al*., 2023). The precise mechanisms by which copper damages bacteria still remain under debate, but several modes of action have been proposed.

One of the main copper toxicity mechanisms is thought to be mis-metallation of proteins by replacing essential metals such as Fe, Mn and Zn, thereby rendering enzymes nonfunctional (Barwinska-Sendra and Waldron, 2017). This includes the disruption of iron-sulfur cluster-containing proteins (Macomber and Imlay, 2009; Chillappagari *et al*., 2010), which are used in a broad range of cellular processes such as central metabolism, respiration and DNA replication. Another proposed mechanism is the generation of reactive oxygen species (ROS) through redox cycling between Cu^+^ and Cu^2+^ in the presence of hydrogen peroxide. Although doubted by some authors (Macomber *et al*., 2007; Saenkham *et al*., 2020; Casanova-Hampton *et al*., 2021), this type of cycling between the two copper oxidation states can generate highly reactive free radicals, leading to oxidative damage of lipids, proteins, and DNA (Sies, 1993; Yoshida *et al*., 1993; Vieira *et al*., 1997; Nan *et al*., 2008). In addition to reactive oxygen species generated by ionic copper, the physical interaction of copper surfaces or nanoparticles with bacterial membranes can also cause oxidative damage to membrane phospholipids and ultimately compromise membrane integrity, leading to cell lysis and death (Santo *et al*., 2008; Hong *et al*., 2012).

Given the toxicity of copper, bacteria must employ efficient defense strategies to survive in the presence of this metal. Under normal physiological conditions, the level of free copper in the cytoplasm is kept extremely low (Outten *et al*., 2000). In Gram-negative bacteria, copper detoxification occurs primarily in the periplasm, where most excess intracellular copper is directed (Changela *et al*., 2003; Macomber *et al*., 2007; Parmar *et al*., 2018). During copper stress, the transcriptional regulator CueR senses cytoplasmic Cu^+^ and activates expression of the copper-export and detoxification genes *copA* and *cueO* (Outten *et al*., 2000; Yamamoto and Ishihama, 2005). CopA is a P-type ATPase that pumps Cu^+^ from the cytoplasm to the periplasm, where the multicopper oxidase CueO oxidizes Cu^+^ into the less reactive and less membrane-permeable Cu^2+^ (Djoko *et al*., 2010). Simultaneously, the CusS/R two-component system detects periplasmic Cu^+^ and induces the CusCFBA efflux pump, which expels copper from the periplasm to the extracellular environment (Rensing and Grass, 2003; Yamamoto and Ishihama, 2005). The periplasmic space in *E. coli* also harbors several proteins that either bind metals or facilitate their transport. Recent research has highlighted the role of periplasmic methionine sulfoxide reductases (MsrPQ), also induced by CusS/R, which repair oxidized methionine residues under copper stress (Vergnes *et al*., 2022). This repair mechanism contributes to the overall cellular health and function during copper exposure, indicating that *E. coli* uses multiple strategies to cope with metal-induced stress.

Copper-induced transcriptional responses in *E. coli* have been characterized previously using DNA microarray analyses. Yamamoto and Ishihama (2004) identified 28 genes that were markedly upregulated and one gene that was downregulated following copper exposure. Their study reported the induction of genes associated with copper homeostasis (*cusB, cusC, cusF, yedV, yedW, copA, cueO*), molybdenum cofactor biosynthesis (*moaB, moaC, moaD, moaE*), and envelope stress and protein quality control (*cpxP, ebr, spy, yccA, ycfS, yqjA, yebE, htpX*), along with several additional genes that could not be clearly assigned to functional categories. A parallel study by Kershaw *et al*. (2005) observed more than a twofold upregulation of 34 genes and the downregulation of 57 genes when comparing a near-lethal concentration (2 mM Cu-glycine) to a well-tolerated level (0,75 mM). According to their results, copper stress induced key copper efflux genes (*copA, cusF*), oxidative stress genes belonging to the SoxRS regulon (*soxS, fldA, fur, zwf*) and envelope stress response genes controlled by the CpxAR system (*cpxP, degP, ompC, ppiD, yccA*). Additionally, many genes involved in iron uptake, storage, or Fe–S cluster assembly (*fiu, cirA, exbD, fecI, fepA, fepB, fes, fhuF, tonB, yecI, ydiC, ydiE, ynh*) were upregulated, including enterobactin biosynthesis genes *(entA, entB, entC, entD, entF)*. The remaining upregulated genes did not cluster into well-defined pathways. Kershaw *et al*. also reported that most of the 57 downregulated genes encoded proteins involved in motility and respiration. Although informative, these DNA microarray-based studies identified only a modest number of copper-responsive genes, likely reflecting the limitations of transcriptomic technologies used in those studies. We therefore sought to revisit copper-induced transcriptional remodeling using RNA sequencing to achieve a more comprehensive, genome-wide view.

To gain genome-wide insight into the transcriptional response to copper, we exposed mid-log-phase *E. coli* cells to both growth-slowing and near-lethal copper concentrations and performed RNA sequencing. To validate the RNA-seq data, we tested the possibility to use the Horizon Discovery *E. coli* genome-wide promoter collection with approximately 1900 promoter-GFP fusions.

## Material and methods

### Media and growth conditions

Bacterial strains were stored as 50% glycerol stocks at −80 °C and streaked onto LB agar plates containing the appropriate antibiotic(s) where applicable. Precultures were grown overnight in 5 mL of medium in 50 mL culture tubes at 37 °C with orbital shaking at 150 rpm. Throughout this paper we use MOPS medium (0.4 M potassium MOPS, 40 mM Tricine, 100 µM FeSO_4_, 95 mM NH_4_Cl, 2.76 mM K_2_SO_4_, 5 µM CaCl_2_, 5.28 mM MgCl_2_, 500 mM NaCl, 30 nM (NH_4_)_6_Mo_7_O_24_, 4 µM H_3_BO_3_, 300 nM CoCl_2_, 100 nM CuSO_4_, 800 nM MnCl_2_, 100 nM ZnSO_4_ (Goormaghtigh and Van Melderen, 2016)) supplemented with 0.4% glucose, 0.4% casamino acids and 0.02 mg/mL tryptophan). For experiments involving the copper biosensor strain, the medium was supplemented with double the amount of glucose, casamino acids and tryptophan (see below). For the Horizon Discovery *E. coli* promoter collection assay and for pH-measurements, LB medium (1% tryptone, 0.5% yeast extract, 1% NaCl (Bertani, 1951)) was used in parallel to MOPS.

### Analysis of CopA induction using copper biosensor

*E. coli* MG1655 harboring two plasmids, a high-copy plasmid pSLcueR (ampicillin 100 µg/mL), encoding the copper-responsive transcriptional regulator CueR, and a medium-copy plasmid pDNP*copA*::*lux* (tetracycline 10 µg/mL), in which the *copA* promoter is transcriptionally fused to the *luxCDABE* operon, was used. In this strain, bioluminescence is induced by bioavailable copper binding to CueR regulatory protein. The system has been described earlier in (Ivask *et al*., 2009) and in this study the original host strain *E. coli* MC1061 was exchanged to MG1655.

The copper biosensor was grown in MOPS medium (5 mL in 50 mL tubes, 150 rpm) supplemented with double the standard concentrations of glucose, casamino acids, and tryptophan (0.8%, 0.8%, and 0.04 mg/mL, respectively) as well as selective antibiotics (10 µg/mL tetracycline and 50 µg/mL ampicillin). This enriched medium was used to improve growth, as the strain exhibits poor growth in standard MOPS due to the combined burden of the high-copy plasmid pSLcueR and dual antibiotic selection. Overnight cultures were diluted to an OD_600_ of 0.05 in fresh medium and 140 µL of the culture was dispensed into each well of a black, clear-bottom 96-well plate (BRANDplates, catalog no. 781671). The culture was then grown to mid-logarithmic phase (OD_600_ ≈ 0.4) in a BioTek Synergy H1 plate reader (Agilent) and subsequently treated with 10 µL of serial dilutions of CuSO_4_ (Biotop) prepared in water, or water alone for the negative control. OD_600_ and luminescence were recorded every 5 min after copper addition.

Statistical analysis was performed using a two-way ANOVA followed by Dunnett’s multiple comparisons test, with correction for multiple comparisons, comparing the control group to each copper concentration at each time point individually.

### Determination of bacterial growth inhibition by copper

*E. coli* MG1655 overnight cultures were diluted to an OD_600_ of 0.05, and 140 µL was transferred to clear 96-well plates unless otherwise specified. Bacteria were grown at 37°C in a BioTek Synergy H1 plate reader (Agilent) with double-orbital shaking at 282 rpm (3 mm orbit) to an OD_600_ of 0.4. For copper treatment, 10 µL of different concentrations of CuSO_4_ (Biotop) solution or water was added and cultures were incubated at 37 °C with shaking. OD_600_ was recorded every 10 minutes.

### Analysis of copper-induced transcriptional effects

#### Bacterial exposure to copper for RNA sequencing

*E. coli* MG1655 (a plasmid-free bacterium from Horizon Discovery *E. coli* promoter collection, see below) was grown overnight in MOPS medium (5 mL in 50 mL tubes, 150 rpm). Then, the culture was diluted to an OD_600_ of 0.05, and 140 µL was transferred to each well of a clear 96-well plate. Cells were grown at 37 °C with shaking in a plate reader to an OD_600_ of 0.4. At OD_600_ = 0.4, 10 µL of water (control) or CuSO_4_ (Biotop) was added to final concentrations of 2 or 8 mM, and cultures were further incubated for 30 min at 37 °C with shaking. 1200 µL of control, 2, or 8 mM CuSO_4_ exposed culture was pooled from microplate wells for RNA extraction and immediately stabilized using RNA protect (Qiagen). Samples were stored at −20 °C for up to one week prior to RNA extraction using RNeasy Mini Kit (Qiagen). RNA sequencing was performed by Novogene, including RNA quality control (RIN measurement), DNase I treatment, rRNA depletion, and library preparation. Sequencing output was ≥2 Gb per sample, with ≥85% of bases achieving Q30 quality. Reads were aligned to the reference genome, and gene expression quantification and differential expression analysis were performed by Novogene. Differential expression was defined as P_adj_ < 0.01 and a ≥2fold change. The data has been deposited in ENA under accession number PRJEB108196.

#### Functional Enrichment Analyses

Gene Ontology (GO) biological process enrichment analysis was performed using the ShinyGO web application (http://bioinformatics.sdstate.edu/go/,version 0.85.1) (Ge *et al*., 2020) with the following parameters: source *EnsemblBacteria* (*Escherichia coli* K-12 MG1655 genes GCF_000005845.2), minimal pathway size: 5 and redundancy removal enabled. To assess GO term overrepresentation, sets of upregulated and downregulated transcripts were compared against the background of all detected transcripts. The statistical analysis is built into the ShinyGO application using false discovery rate cutoff of p < 0.001.

### Horizon Discovery E. coli promoter collection assay

The *E. coli* promoter library was obtained from Horizon Discovery (https://horizondiscovery.com/en/non-mammalian-research-tools/products/e-coli-promoter-collection). This genome-scale collection consists of fluorescent reporter strains in which individual *E. coli* K-12 MG1655 promoters are fused to a bright, fast-folding GFPmut2 on a low-copy plasmid referred to as pUA66 in academic literature (Zaslaver *et al*., 2006) and pMSs201 commercially. The collection includes over 1,900 promoters to enable high-resolution monitoring of promoter activity across the genome. Promoters were defined as intergenic regions longer than 40 bp, and during library construction 50–150 bp from each flanking gene were included upstream of *gfp* to preserve native regulatory context (Zaslaver *et al*., 2006). Selected strains from the Horizon Discovery *E. coli* promoter collection – corresponding to the 12 most strongly up- and downregulated genes identified in our RNA-seq analysis–were sequenced using the pUA66-fw primer (CAACCTTACCAGAGGGCG). Two genes, *tnaC* and *ybcV*, were excluded from further testing due to unsuccessful sequence confirmation for the *E. coli* promoter collection construct. The promoter collection strains were grown in MOPS medium supplemented with 0.4% glucose, 0.4% casamino acids and 0.02 mg/mL tryptophan or LB medium with 25 µg/mL kanamycin. OD_600_ and GFP fluorescence (excitation: 485 nm, emission: 528 nm) were recorded every 10 minutes. Background was subtracted from fluorescence and OD600 measurements. OD_600_-normalized fluorescence of Cu-treated cultures relative to the no-copper control was calculated for each timepoint:

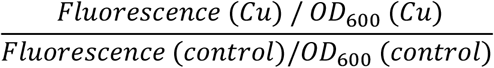

### Measurement of growth-media pH

*E. coli* MG1655 was grown in 10 mL MOPS or LB medium at 37 °C until OD_600_ ≈ 0.4. Cultures were then treated with CuSO_4_ to final concentrations of 2 mM or 8 mM for 30 min, pelleted at 4,430 × g for 7 min, and the supernatants were passed through 0.2 µm Minisart syringe filters (Sartorius). pH was measured with SevenDirect SD20 (Mettler Toledo).

Statistical analysis was performed using two-way ANOVA followed by Dunnett’s multiple comparisons test, comparing untreated controls to each copper concentration within each medium.

### GFP quenching experiment

*E. coli* MG1655 was grown overnight in a plate reader as explained in “Determination of bacterial growth inhibition by copper” for 18 hours. The stationary phase bacteria were challenged with different concentrations of CuSO_4_, AgNO_3_ and benzalkonium chloride. OD_600_ and GFP fluorescence (excitation: 485 nm, emission: 528 nm) were recorded every 2 minutes.

## Results

### Low millimolar copper affects E. coli within minutes

To determine the growth conditions for investigating the *E. coli* copper stress response in MOPS medium, we conducted a preliminary minimal inhibitory concentration (MIC) assay. We found that *E. coli* tolerates 2-8 millimolar copper with increasing difficulty and 16 mM copper already inhibits growth completely (Supplementary Figure 1). Therefore, we chose 2 mM CuSO_4_ as a concentration that slows growth, and 8 mM CuSO_4_ as a near-lethal level that only allows initial growth before cell numbers dwindle. Growth curves of the actual cultures used for RNA extraction grown with 2 and 8 mM Cu are shown in Figure 1 B.

**Figure 1.**
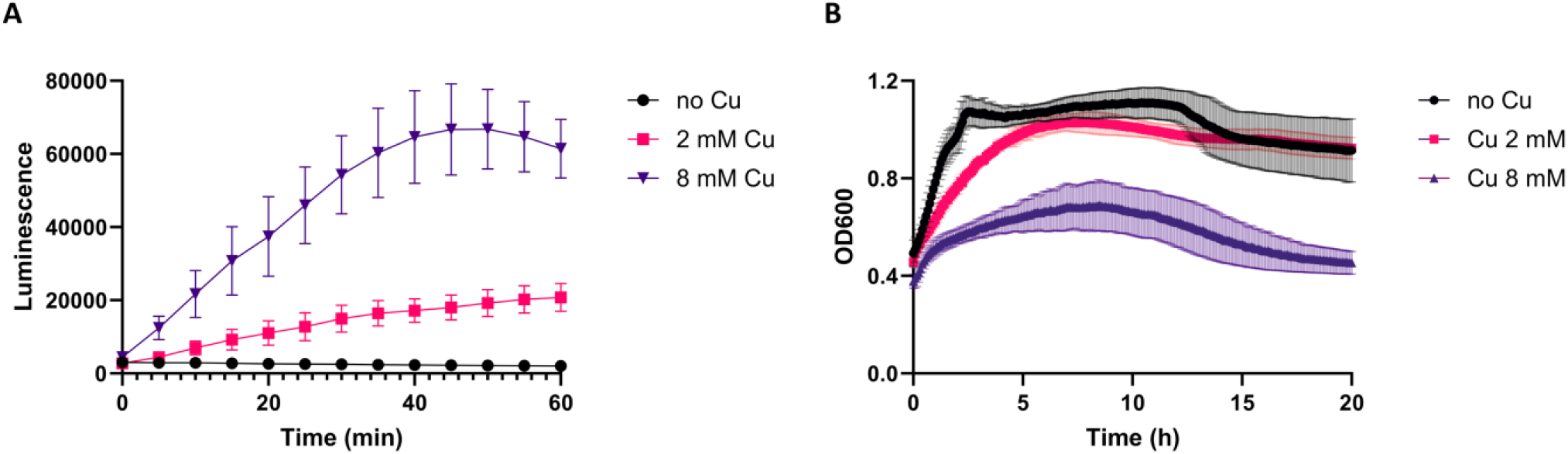
**A. Determining an appropriate copper exposure time using a bioluminescent biosensor**. *E. coli* MG1655 carrying pSLcueR and pDNPcopA::lux (Ivask *et al*., 2009) responding to 2 and 8 mM CuSO_4_ over time. Means ± 95% confidence intervals (CI) of two independent experiments with seven technical replicates each are shown. **B. Growth of cultures used for RNA-seq**. Growth curves of *E. coli* MG1655 in MOPS with 2 and 8 mM CuSO_4_. Cultures shown are the same ones sampled at 30 min for RNA extraction. The error bars represent mean ± 95% CI of three independent experiments with one to two replicates.

As previous transcriptional analyses have used widely different copper treatment times - 5 minutes (Yamamoto and Ishihama, 2005) and 10 hours (Kershaw *et al*., 2005) - we then sought to determine an appropriate treatment duration for our RNA-sequencing experiments. To do this, we used a copper-responsive biosensor (Ivask *et al*., 2009) that produces increased bioluminescence in the presence of elevated intracellular copper. Exposure to 8 mM CuSO_4_ caused a significant rise in bioluminescence as early as 5 minutes (P_adj_ < 0.0001), while 2 mM CuSO_4_ produced a significant increase at 20 minutes (P_adj_ < 0.0001) (Figure 1 A). Both concentrations continued to induce an increasing signal over time. Based on these results, we selected 30 minutes as the treatment duration for the subsequent RNA-sequencing experiment.

### Copper exposure upregulates oxidative stress response, histidine production and iron acquisition genes

To investigate how *E. coli* MG1655 responds to copper stress, we performed transcriptional profiling of exponentially growing cells in MOPS medium treated with growth-inhibitory 2 mM copper and near-lethal 8 mM copper (Figure 1 B). With a cutoff of P_adj_ < 0.01 and a minimum 2-fold change, 2 mM copper **upregulated 487 genes**, whereas 8 mM Cu **upregulated 364 genes**, with an overlap of 296 genes (Supplementary file 2). This confirmed that the chosen Cu treatment time was relevant and sufficient to induce transcriptomic rearrangements.

To identify biological functions affected by copper stress, we performed Gene Ontology (GO) analysis to determine which terms were enriched in the Cu-exposed bacteria. Treatment with 8 mM Cu resulted in 13 enriched GO terms, while 2 mM Cu yielded 19 terms, including all those found at 8 mM (Figure 2 A, B, Supplementary file 1). Categories enriched at both Cu concentrations included response to copper ion, response to oxidative stress, histidine biosynthetic process, and multiple GO terms related to iron acquisition, such as *E. coli* siderophore (enterobactin) biosynthesis. Additional terms enriched only at 2 mM Cu were response to heat and protein folding as well as biosynthetic processes for methionine, arginine and lipid A (Figure 2 A, Supplementary file 1). Below, we describe in detail the genes assigned to each of the major upregulated functional categories.

**Figure 2.**
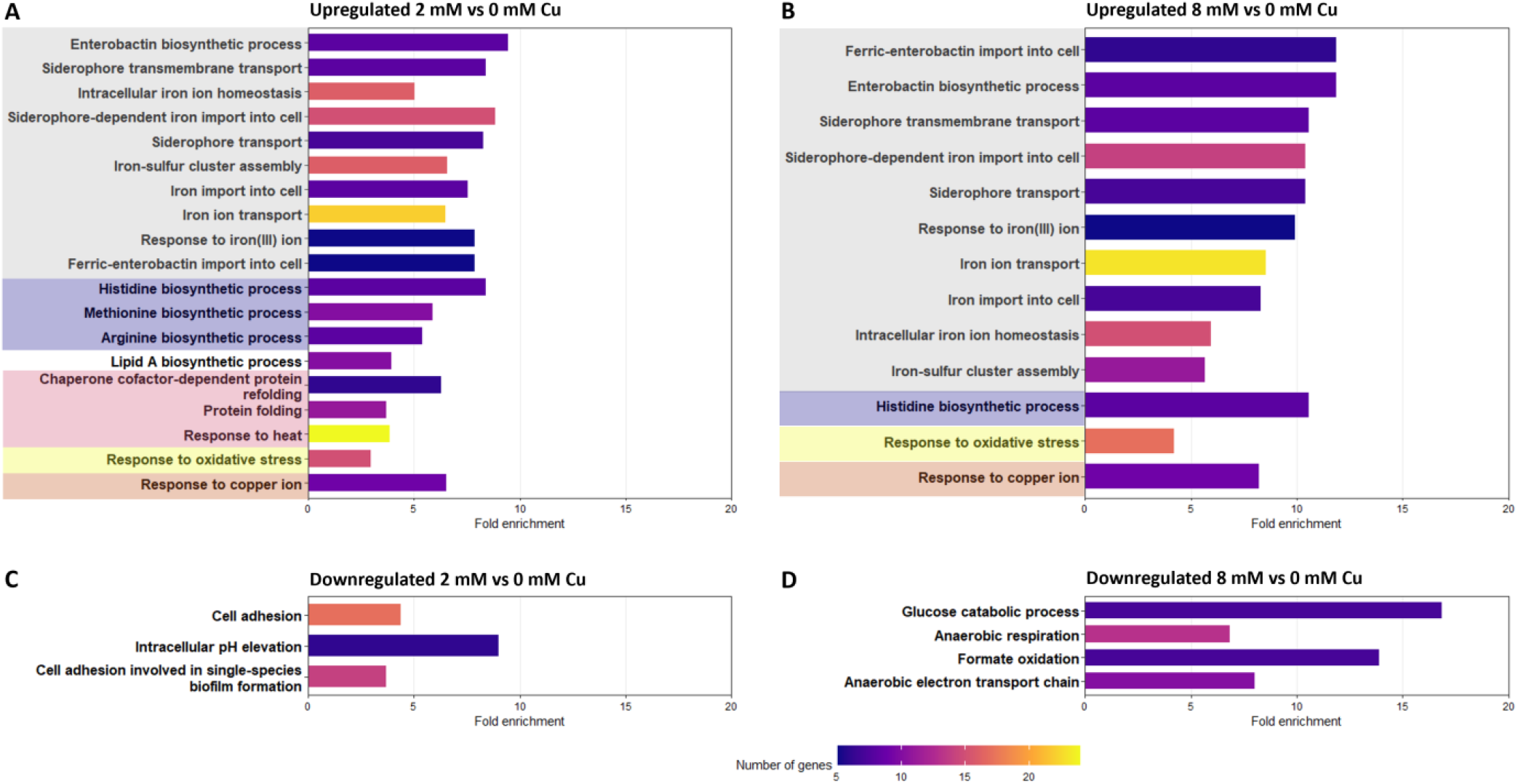
Transcriptional profiling of *E. coli* under copper stress. GO enrichment analysis of *E. coli* MG1655 genes upregulated at 2 mM Cu (**A**) and 8 mM Cu (**B**) and downregulated at 2 mM Cu (**C**) and 8 mM Cu (**D**) compared to no Cu control. Only statistically significantly up- or downregulated GO terms that contain at least 5 genes are shown.

As expected, several **known copper responsive genes** were upregulated upon elevated Cu exposure. Exposure to 2 mM Cu upregulated 13 genes associated with copper stress response and homeostasis. Of these, nine were identified through GO enrichment analysis (Figure 2, orange background, Supplementary file 1), and four additional genes (indicated with an asterisk) were identified by manual inspection of the differentially expressed gene set (supplementary file 2). The identified genes are responsible for transporting excess copper out (*copA*^*\**^, *zntA*^*\**^, *cusA, cusB, cusC, cusF*), regulating the copper response (*cusS*^*\**^*/cusR, yedV/yedW*^*^), detoxifying copper in the periplasm (*cueO*) and possibly trying to keep copper from entering the bacterium (*bhsA*). Curiously the expression of *comR* (*ycfQ*), the repressor for *bhsA*, has gone up at the same time as *bshA* itself. The expression of *comR* is only modestly upregulated, 2.24-fold at 2 mM Cu and 2.02-fold at 8 mM Cu, whereas *bshA* shows much stronger induction, increasing 13.79-fold at 2 mM Cu and 10.91-fold at 8 mM Cu. Therefore, it is possible that the slight increase in *comR* expression is due to random fluctuation and not a true effect.

Copper stress strongly induces the **iron acquisition response genes** (Figure 2, grey background, Supplementary file 1). In total, 59 iron-related genes are upregulated in the presence of 2 mM Cu, and 55 genes show upregulation at 8 mM Cu (Supplementary file 1). This response includes components involved in the uptake of both ferrous (Fe^2+^) and ferric iron (Fe^3+^). The whole feo system, responsible for Fe^2+^ uptake (*feoA, feoB, feoC*), is induced. Fe^3+^ acquisition pathways are more complex, yet also strongly upregulated during copper exposure. This includes the induction of the full fec operon (*fecA, fecB, fecC, fecD, fecI, fecR*), mediating ferric-citrate transport, as well as enterobactin biosynthesis genes (*entA, entB, entC, entD, entE, entF, entH, entS, ybdZ*) and their corresponding transport and processing genes (*fepA, fepB, fepC, fepD, fepG, fes, fhuF*).

GO enrichment analysis also revealed induction of **amino acid biosynthesis genes** in copper-exposed bacteria (Figure 2, violet background, Supplementary file 1). Histidine biosynthesis genes (*hisF, hisA, hisH, hisI, hisG, hisC, hisB, hisD*) were enriched at both copper concentrations, while methionine (*metA, metF, metH, metE, mmuP, metC, metL, metB, metR, thrA*) and arginine (*argE, argI, argH, argF, argA, argB, argC, argD*) biosynthesis genes were additionally enriched at 2 mM Cu.

Exposure to both 2 mM and 8 mM Cu resulted in a marked enrichment of **oxidative stress response genes** (Figure 2 yellow background, Supplementary file 1). A total of 12 genes were upregulated at both copper concentrations, while an additional 3 genes were induced exclusively at 2 mM Cu and 5 genes exclusively at 8 mM Cu. Collectively, they are linked with protection against peroxides (*oxyR, katG, ahpC, ahpF, hslO, degP, azoR*), defence against superoxide O_2_^-^ (*soxR, soda, ydbK, ytfK*) and Fe-S cluster repair (*sufA, sufD*, *sufE, nfuA, fumC, acnA*). The prominence of Fe-S cluster damage under Cu stress is further supported by the strong enrichment of the Gene Ontology term “Fe-S cluster assembly,” encompassing 16 genes involved in Fe-S cluster biogenesis.

Three GO terms associated with **heat response and protein folding**: “response to heat”, “protein folding” and “chaperone cofactor-dependent protein refolding” are significantly enriched at 2 mM Cu, encompassing 27 unique genes (Figure 2, pink background, Supplementary file 1). Bulk of these genes encode proteins that reverse or prevent protein misfolding (*dnaK, dnaJ, grpE, groL, groS, hscA, hscB, clpB, ibpA, ibpB, htpG, hslO, ppiA*) or degrade misfolded proteins (*clpP, hslU, hslV, lon, degP, hflK, hflC*). A smaller subset represents general or heat-induced stress-response genes (*rpoH, rpoD, marR, pspA, sodA, moaA, hslR*). Although *rpoD* functions primarily as the housekeeping sigma factor, it is known to be upregulated under heat-stress conditions (Park *et al*., 2024).

Exposure to 2 mM Cu resulted in a 3.9-times enrichment of **lipid A biosynthesis genes** (Figure 2 A, Supplementary file 1). Lipid A is the molecule that anchors the lipopolysaccharide molecule into the outer membrane. In total 10 lipid A biosynthesis genes (*arnT, arnD, arnC, arnE, eptA, lpxC, arnB, arnA, eptB*) are upregulated.

### Copper exposure represses genes related to adhesion and anaerobic respiration

Exposure to 2 mM Cu **downregulated 486 genes** while 8 mM Cu **repressed 217 genes**, with an overlap of 129 genes (Supplementary file 2). Interestingly the enriched GO terms for downregulated genes between 2 and 8 mM Cu do not overlap.

Exposure to 2 mM Cu downregulated 19 genes involved in bacterial **adhesion to surfaces** belonging to GO categories “cell adhesion” and “cell adhesion involved in single-species biofilm formation” (Figure 2 C, Supplementary file 1). Most of these genes encode confirmed or predicted adhesins (*fimH, elfA, elfG, sfmA, yadC, yadL, yadM, ybgO, ycgV, ygiL, ypjA, yqiI*). Additional downregulated genes include type 1 fimbriae genes (*fimA, fimF, fimG, fimI*) known to be important for attachment and biofilm formation (Lasaro *et al*., 2009). The remaining three genes encode a flagellar biofilm exopolysaccaride PGA synthase *pgaD*, filament capping protein *fliD*, and putative c-di-GMP binding protein *cdgI*.

Unexpectedly, the enrichment analysis revealed that genes involved in **intracellular pH elevation** (*gadB, adiA, gadC, gadA, kefC, kefF*) were downregulated in response to 2 mM Cu (Figure 2 C, Supplementary file 1). Since copper acidifies the growth medium (Gunawan *et al*., 2011; Figure 6), we would expect these pH-regulating genes to be upregulated, not repressed.

Exposure to 8 mM Cu downregulated 12 genes associated with **anaerobic energy metabolism**, falling into the GO categories “anaerobic respiration,” “glucose catabolic process,” “formate oxidation,” and “anaerobic electron transport chain” (Figure 2 D, Supplementary file 1). These include genes encoding subunits of hydrogenase 1 (*hyaA, hyaC*) which mediates H_2_ oxidation, hydrogenase 2 (*hybA, hybB, hybO*), which mediates hydrogen oxidation and production (Laurinavichene and Tsygankov, 2001; Lukey *et al*., 2010), as well as hydrogenase 3 (*hycB, hycC, hycD, hycF, hycG*), responsible for H_2_ generation from formate as part of the formate hydrogenlyase complex (Maeda *et al*., 2007). Additional downregulated genes include *fdhF*, which provides electrons to hydrogenase 3, and *dcuB*, encoding the fumarate/succinate antiporter required for fumarate respiration.

### Cu-stressed E. coli are not relieved by external iron supplementation

Because our RNA-seq data showed strong induction of iron-acquisition genes during copper stress, we considered the possibility that the cells were experiencing functional iron limitation. To test whether supplemental iron could alleviate copper-induced stress, we measured the copper MIC in the presence of 100 or 200 µM FeSO_4_ (Figure 3). The addition of FeSO_4_ did not improve the growth of copper-treated cells, indicating that external iron does not relieve the effects of copper stress under these conditions.

**Figure 3.**
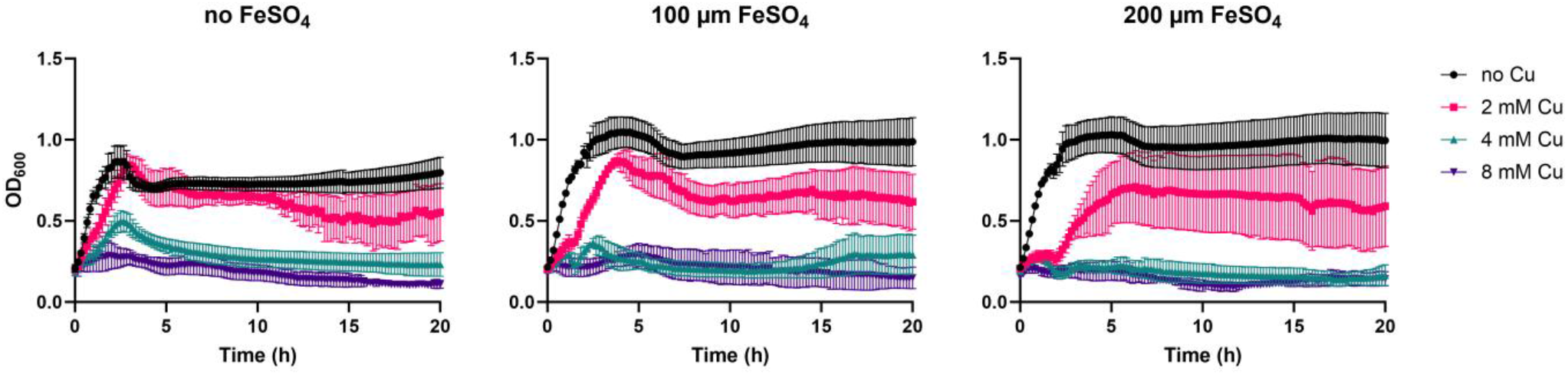
Influence of iron supplementation on copper-induced growth inhibition. Growth curves of *E. coli* MG1655 in MOPS medium supplemented with 2, 4 or 8 mM CuSO_4_, with or without added FeSO_4_. The errors bars represent mean ± 95% CI of three independent experiments with three replicates.

### Limited utility of the Horizon Discovery E. coli promoter collection for screening copper-responsive promoters

Before measuring the effect of copper on the transcriptome via RNA sequencing, we attempted to study cellular response to copper exposure using a genome-wide *E. coli* promoter collection from Horizon Discovery. This library contains 1,900 promoters from *E. coli* MG1655, each individually cloned upstream of a fast-folding GFP in a kanamycin-resistant plasmid (pMS201, also referred to as pUA66 in the literature, (Zaslaver *et al*., 2006)). This setup enables screening for genes that are up- or downregulated when treated with a chemical by measuring the GFP fluorescence in strains carrying different promoters. When the library strains were exposed to 1 mM copper, the signal-to-noise ratio was extremely low (data not shown), making this approach unsuitable for a reliable primary screening. The same issue of low signal to noise ratio was observed also when an attempt was made to study the response of the promoter library to 12.5 µM silver and 2.5 µM benzalkonium chloride (data not shown) referring to the fact that the low signal issue was not specifically related to copper exposure. We therefore explored whether the promoter collection could be used to validate RNA-seq results. From the RNA-seq dataset, we selected 12 genes that were among the most strongly upregulated or downregulated and were present in the promoter collection. The 24 promoter regions in the library were verified by sequencing and two downregulated genes were excluded due to unsuccessful sequence confirmation. Copper concentrations of 1 mM and 0.5 mM were chosen for this experiment as the highest concentrations that did not affect growth of MG1655 carrying the promoter collection library plasmid pUA66 (Supplementary Figure 2). MOPS medium was used to ensure consistency with the RNA-seq experiments, whereas LB medium was selected because it is the manufacturer-recommended growth medium for the Horizon Discovery *E. coli* promoter collection.

Of the 12 promoters highly upregulated in the RNA-seq dataset, only two exhibited more than a twofold increase in GFP fluorescence (log_2_fold change (log_2_FC) > 1) compared to the control without copper. The *cusC* promoter-GFP fusion showed the strongest response among the tested strains and was induced by copper in both used media, LB and MOPS medium (Figure 4). Induction of this promoter constructs started within the first minutes of copper exposure and reached its maximum after 30 min. In the presence of 0.5 mM and 1 mM of copper, fluorescence increased on average 2.9-fold (log_2_FC = 1.5) and 2.5-fold (log_2_FC = 1.3), respectively. In LB medium those copper concentrations induced the fluorescence by 5.6- and 6.0-fold (log_2_FC = 2.5 and 2.6), respectively. Considering that the *cusS* gene showed a dramatic 2293-fold (log_2_FC = 11.2, Supplementary Figure 1) upregulation in RNA-seq assay, the promoter construct upregulation was truly modest. For the *yebE* promoter-GFP fusion, no induction was observed in LB medium and in MOPS medium only a modest 2.1-fold increase (log_2_FC = 1.1) was observed (Figure 4). Interestingly, in the presence of 1 mM Cu in MOPS medium, no induction was observed. Analogously to *cusS*, also *yebE* was highly upregulated in the RNA-seq assay with 135-fold (log_2_FC = 7.1) induction. Moreover, the rest of the 10 promoters selected due to their upregulation in RNA-seq assay did not seem to be activated at all (Figure 4).

**Figure 4.**
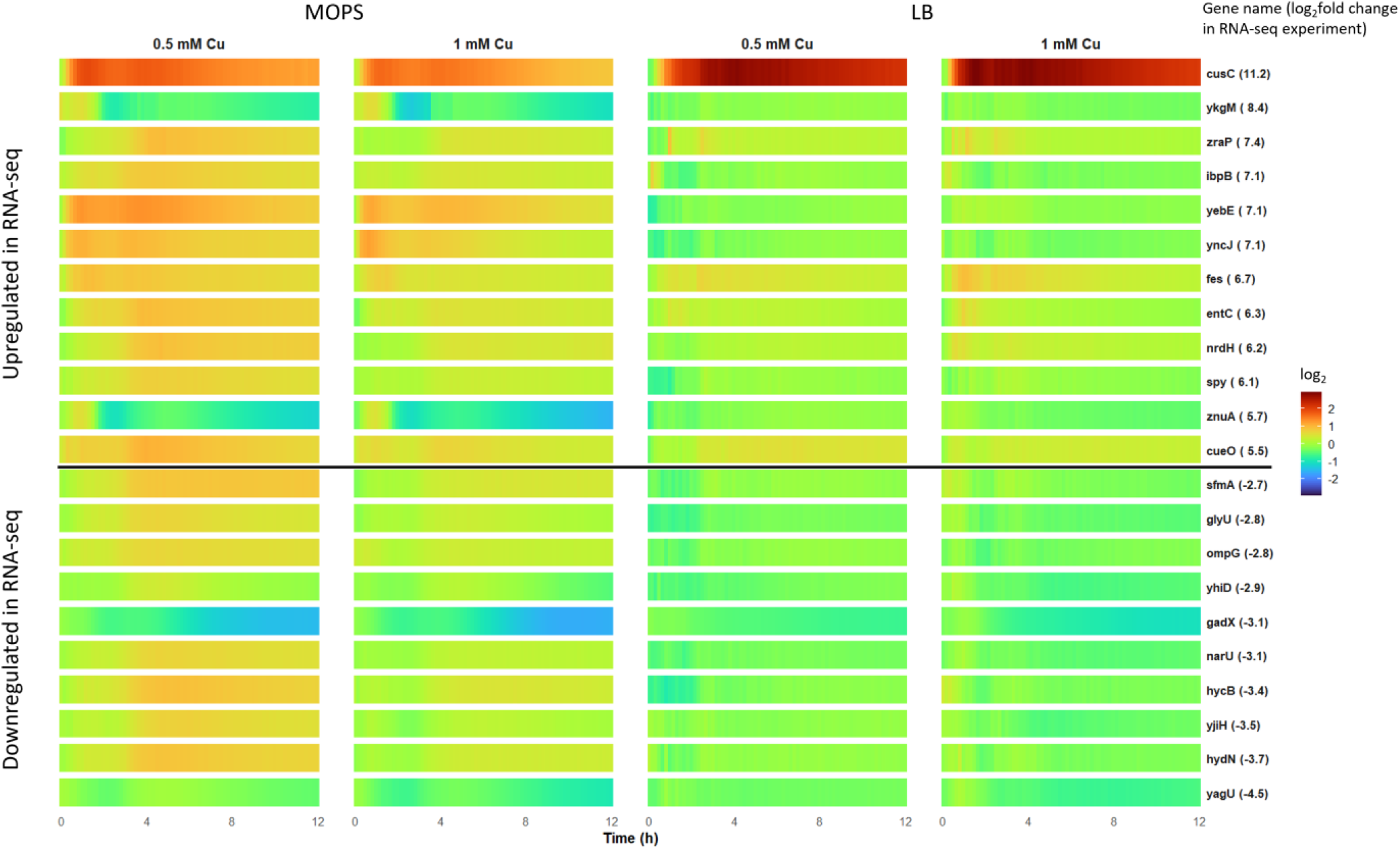
Using Horizon Discovery E. coli promoter collection to study gene expression changes due to copper stress. Dynamics of OD_600_-normalized fluorescence for GFP-based Horizon Discovery *E. coli* promoter collection strains relative to the no-copper control over time (log_2_ scale). Red tones indicate fluorescence activation, with deeper shades reflecting stronger induction, while green represents no change and blue tones denote inhibition. Data represent means of four independent experiments in LB and five in MOPS.

Among the 10 promoters strongly downregulated in RNA-seq, only *gadX* consistently showed repression in the GFP-based assay (Figure 4). Excluding the first 6 hours, GFP signal from the *gadX* promoter decreased on average by 2.5-fold (log_2_FC = −1.3) with 0.5 mM Cu and 2.7-fold (log_2_FC = −1.4) with 1 mM Cu in MOPS. In LB, downregulation was less pronounced, remaining below twofold.

In conclusion, of the 12 promoters identified as highly upregulated by RNA-seq, only two (*cusC* and *yebE*) showed any measurable activation, and in both cases the response was modest, with *yebE* exhibiting only borderline induction. Similarly, among the 10 promoters classified as downregulated in the RNA-seq dataset, only *gadX* displayed consistent repression in the reporter assay. These results indicate that the Horizon Discovery *E. coli* promoter collection is not well suited for assessing copper-dependent transcriptional responses under the conditions tested.

One possible explanation for the weak performance of the promoter collection in copper-exposure assays is fluorescence quenching. Both Cu^+^ and Cu^2+^ ions are known to strongly quench ST-Kaede and ST-GFP fluorescent proteins, resulting in near-complete loss of fluorescence at around 0.4 mM CuSO_4_ (Bourge *et al*., 2015). To assess whether copper quenches GFP under our experimental conditions, we monitored GFP fluorescence in an *E. coli* strain carrying a promoter library construct with the strong *rpsT* promoter. Cultures were grown to stationary phase and then treated with Cu or one of three controls: H_2_O, Ag, or benzalkonium chloride (BAC). Aside from a brief drop in fluorescence associated with pausing plate-reader shaking, none of the treatments: copper, silver or benzalkonium chloride – caused a substantial decrease in fluorescence relative to the water control (Figure 5). Therefore, the low signal-to-noise ratio observed in the Horizon Discovery *E. coli* promoter collection is unlikely to be attributable to copper-induced quenching of GFP fluorescence.

**Figure 5.**
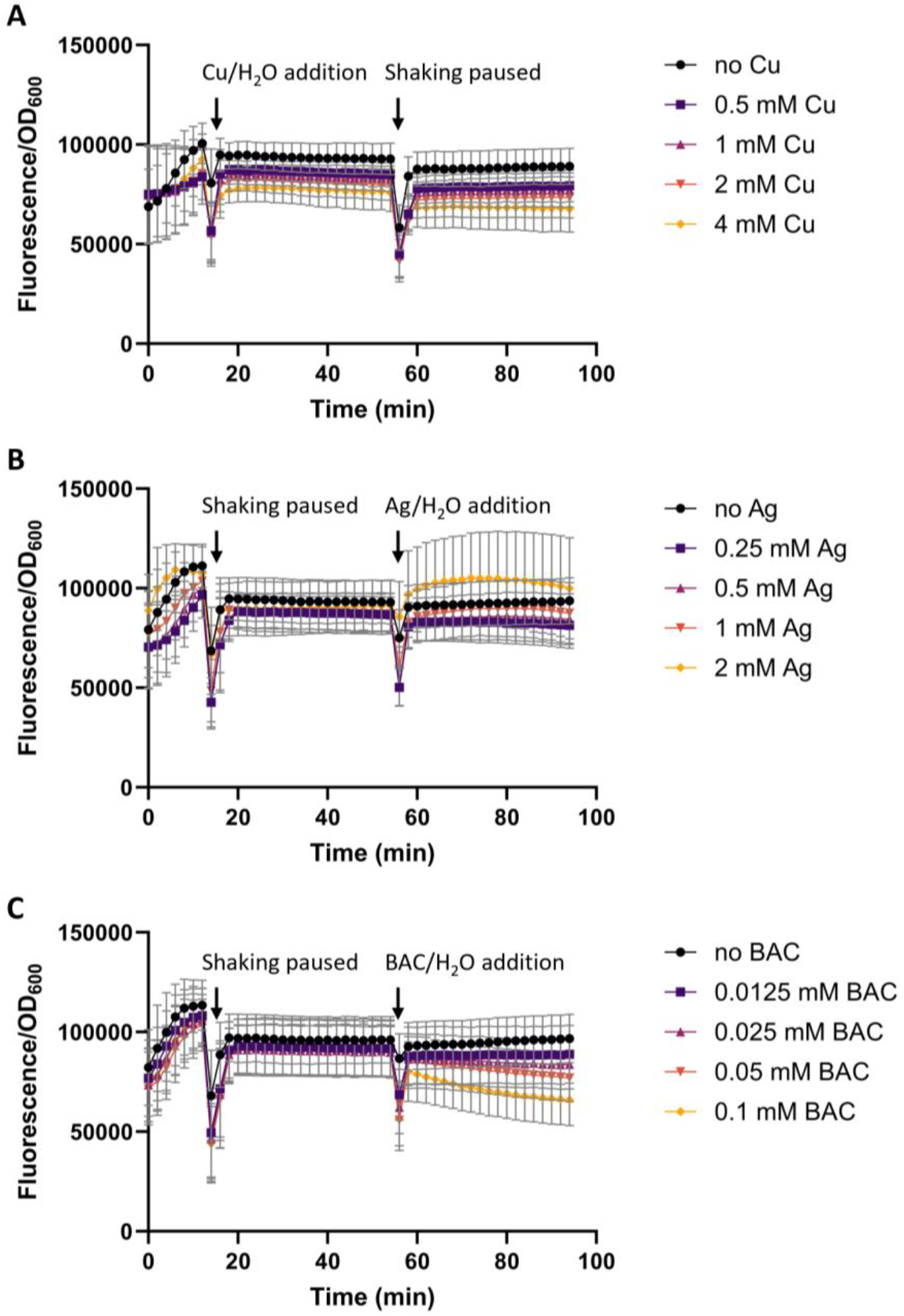
Quenching of GFP signal. OD_600_ normalised fluoresence of Horizon Discovery *E. coli* promoter collection strain MG1655 with pUA66 *rpsT* in the presence of copper (**A**), silver (**B**) or benzalkonium chloride (BAC) (**C**). Means ± 95% CI of four independent experiments with two to three technical replicates each are shown. “Shaking paused” indicates timepoints when the plate reader was briefly stopped to add other chemical(s), while nothing was added to this sample.

### Copper-induced acidification is minimal in MOPS medium

From previous studies it is known that copper may acidify bacterial growth media (Gunawan *et al*., 2011). To assess whether the transcriptomic responses to copper exposure could be instead caused by copper-induced growth media acidification, the pH of *E. coli* cultures after 30 min exposure to 2 and 8 mM copper was measured in the same buffered MOPS medium used for RNA-seq, and LB medium was included as an unbuffered control. In LB, the pH significantly decreased from 6.5 to 5.6 with 2 mM Cu and further to 3.9 with 8 mM Cu (Figure 6). In the buffered MOPS medium it remained more stable: the pH without added Cu was 6.7, in the presence of 2 mM Cu the pH was also 6.7, and in the presence of 8 mM Cu the pH was 6.1.

**Figure 6.**
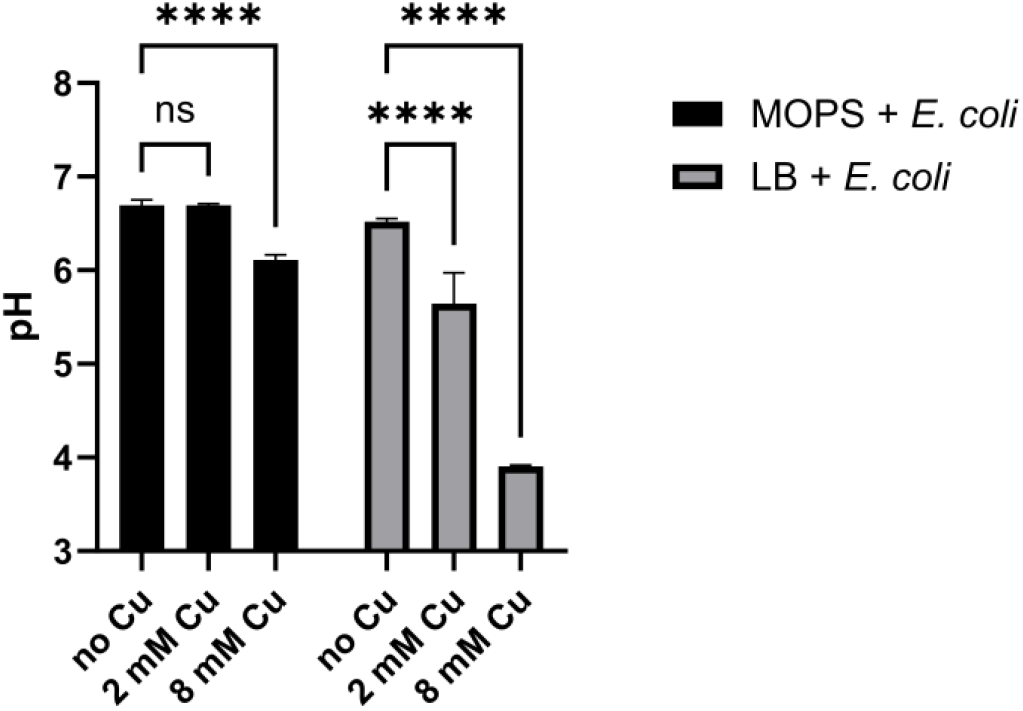
Effect of copper exposure on culture pH. Changes in pH after 30-min copper exposure in *E. coli* cultures grown in MOPS or LB medium. Means ± 95% CI of three independent experiments with three technical replicates each are shown.

## Discussion

Our RNA-seq analysis showed that copper stress triggers a broad and coordinated remodeling of *E. coli* gene expression. This was evident under both concentrations: 2 mM copper, which slows bacterial growth, and 8 mM copper, a near-lethal concentration that only allows limited growth before cell density declines (Figure 1 B; Supplementary Figure 1). Exposure to 2 mM copper elevated the expression of 487 genes and 8 mM copper upregulated 364 genes with a substantial overlap between the two conditions (Supplementary file 2). Firstly, all canonical copper-stress response genes were activated at both copper concentrations (Figure 2, orange background, Supplementary file 1). These include the Cu^+^ exporter CopA, the CusABCF efflux system, the periplasmic multicopper oxidase CueO, and the copper-responsive two-component systems CusS/CusR and YedV/YedW (Supplementary file 1). The induction of all major components of the known copper-response network indicates that our treatment conditions were appropriate for eliciting a robust copper stress response.

Our RNA-sequencing data also revealed upregulation of **histidine, methionine and arginine biosynthesis genes** in response to copper exposure (Figure 2, violet background, Supplementary file 1). It has been previously speculated that elevated histidine biosynthesis may be important in protection against copper toxicity (Sullivan *et al*., 2024) as histidine is a known amino acid to bind both Cu^2+^ and Cu^+^ (Mehlenbacher *et al*., 2015; Brander *et al*., 2020). Here we show that in *E. coli* copper stress upregulates histidine biosynthesis genes with both copper concentrations: growth-slowing 2 mM and near-lethal 8 mM (Figure 2, violet background, Supplementary file 1). Histidine can thereafter bind copper ions, reducing free copper levels and protect the bacterium from copper toxicity. In addition to binding copper, histidine acts also as a buffering agent (Abe, 2000). This could represent another layer of defense against copper stress in unbuffered media where copper lowers the pH dramatically (Figure 6)(Gunawan *et al*., 2011). However, we observe the upregulation of the histidine biosynthesis pathway in in buffered MOPS medium, where the pH reduction is minimal (Figure 6). Therefore, we propose that intracellular Cu chelation is the main benefit of high histidine levels. Moderate 2 mM copper exposure also upregulated methionine and arginine biosynthesis genes (Figure 2, violet background, Supplementary file 1). Methionine biosynthesis is probably upregulated for the same reason as histidine, as methionine can also bind both Cu^2+^ and Cu^+^ (Contaldo *et al*., 2025). Why arginine biosynthesis gets upregulated is less straightforward as it binds copper very weakly (Wu *et al*., 2010; Bottari *et al*., 2019).

In the field of copper toxicity, there is a divide between authors who believe that oxidative stress is an important copper toxicity mechanism and those who believe it not to be. Here we observed that 15 **oxidative stress response genes** are upregulated during moderate copper stress and 12 under high copper stress (Figure 2, yellow background). These include genes involved in protection against peroxides, defense against superoxide O_2_^-^, and Fe-S cluster (Supplementary file 1). Although the upregulation of oxidative stress response genes suggests that oxidative stress is involved in bacterial copper response, this alone does not establish it as the dominant effect of copper exposure but at the same time does not allow it to be discarded either.

Our RNA-seq data indicates that copper stress robustly induces **genes involved in iron acquisition** (Figure 2, grey background, Supplementary file 1). This encompasses genes involved in the uptake of both ferrous Fe^2+^ and ferric Fe^3+^, including those required for enterobactin biosynthesis, processing, and transport. At first glance, this transcriptional profile resembles a state of severe iron starvation. It is known that Cu mis-metallates proteins by replacing Fe (Barwinska-Sendra and Waldron, 2017), which suggests that additional Fe may be needed to produce new enzymes. However, previous studies have shown that upon copper exposure the total measurable Fe does not significantly decrease in *E. coli* cells (Casanova-Hampton *et al*., 2021), and we show here that external iron supplementation does not relieve the copper stress (Figure 3). This suggests that the strong activation of iron acquisition pathways under copper stress does not reflect true iron scarcity.

As mentioned above, we show here that among other iron acquisition genes, many enterobactin biosynthesis, transport and processing genes are upregulated in copper stress (Figure 2 A, B, Supplementary file 1). The link between enterobactin and copper stress has been drawn before, however it is a little complicated. Mutants defective in enterobactin production are more sensitive to Cu (Casanova-Hampton *et al*., 2021; Peralta *et al*., 2022), indicating that enterobactin has a protective role. This has been linked with reducing copper-induced oxidative stress: strains lacking enterobactin are more susceptible to copper-induced oxidative stress, and physiological enterobactin concentrations confer protective effects by reducing ROS levels (Adler *et al*., 2014; Peralta *et al*., 2016; Peralta *et al*., 2022). However, high enterobactin levels may promote Cu^2+^ reduction to Cu^+^ and increase toxicity (Peralta *et al*., 2022). The copper-siderophore chemistry is context-dependent: enterobactin can act as an antioxidant under certain redox conditions and cellular localizations, but it can also participate in redox cycling that enhances copper toxicity if present in excess or not properly processed (Kim *et al*., 2001; Grass *et al*., 2004; Peralta *et al*., 2022).

Under moderate copper stress (2 mM), we observed also **upregulation of heat-shock-associated genes**, which largely overlap with protein folding and refolding pathways (Figure 2, pink background, Supplementary file 1). This suggests that proteins in Cu-stressed cells experience substantial damage or misfolding. Such disruption is expected, given that copper can mis-metallate proteins (Barwinska-Sendra and Waldron, 2017) and destroy Fe-S clusters (Macomber and Imlay, 2009; Chillappagari *et al*., 2010), leaving enzymes inactive and destabilized. Together, these effects likely create a substantial burden of misfolded or damaged proteins, triggering a pronounced heat-shock response.

Under moderate copper stress, we also observed an enrichment of lipid A biosynthesis genes (Figure 2 A, Supplementary file 1), including the key enzyme LpxC (Supplementary file 2). However, since excessive lipid A synthesis is harmful for bacteria (Hummels, 2025), we cannot at this moment distinguish whether the upregulation is a beneficial reaction to, or an unintended consequence of copper stress.

Copper exposure also caused broad gene repression across the transcriptome. At 2 mM Cu, 486 genes were downregulated (Supplementary file 2); however, enrichment analysis only showed enrichment in categories related to biofilm formation and, unexpectedly, pathways linked to pH elevation (Figure 2 C, Supplementary file 1). The reduction in biofilm-related genes fits with the idea that copper-stressed bacteria are less inclined to settle on surfaces, which is consistent with previous reports showing that copper inhibits biofilm formation of *E. coli* and *Listeria monocytogenes* at both MIC and sub-MIC concentrations (Dehkordi *et al*., 2023). The repression of genes involved in pH elevation at 2 mM Cu is more difficult to interpret. Notably, the glutamate decarboxylase genes *gadA* and *gadB* were downregulated. One possible explanation is that copper damages glutamate synthase – a Fe-S-containing enzyme essential for synthesizing glutamate (Djoko *et al*., 2017). Because GadA and GadB convert glutamate into GABA to help raise intracellular pH, continuing to express these enzymes would consume glutamate. If copper stress limits glutamate production, reducing GadAB expression may therefore help the cell conserve its remaining glutamate, especially when the external pH remains relatively stable and the need for GadAB-mediated pH regulation is lower (Figure 6). At 8 mM Cu, 217 genes were repressed, and enrichment analysis showed enrichment only in anaerobic respiration and hydrogenase-based energy metabolism. The repression of anaerobic respiration and hydrogenase-linked pathways at 8 mM Cu is consistent with copper’s known ability to interfere with Fe–S cluster assembly, which both processes depend on (Tan *et al*., 2017).

As an alternative approach to expression profiling, we evaluated the Horizon Discovery promoter library, but it proved unsuitable due to poor signal-to-noise ratios during copper exposure. Of the highly copper-responsive genes identified by RNA-seq, only *cusC* showed strong activation in the GFP-based assay (Figure 4). Another gene, *yebE*, exhibited only a modest expression from its promoter in MOPS at 0.5 mM Cu, and failed to reach the twofold induction threshold in all other conditions. The remaining ten strongly upregulated genes detected by RNA-seq showed no measurable activation in the promoter collection. A similar pattern was observed for copper-repressed genes: of the ten promoters strongly downregulated in RNA-seq, only *gadX* promoter consistently showed repression in MOPS. The other nine promoters displayed no detectable decrease in GFP signal.

The underlying cause of the poor performance of the collection remains unresolved. We considered fluorescence quenching by copper, since in some conditions it practically removes GFP fluorescence (Bourge *et al*., 2015) by causing FRET (Hötzer *et al*., 2011; Jensen *et al*., 2024). Our experiments indicate, however, that under our tested conditions, copper does not affect intracellular GFP fluorescence (Figure 5). In essence, compared to the direct readout of transcriptional activity by RNA-seq, a GFP-based system is more convoluted. Any change in fluorescence levels requires not only transcription but also translation and maturation of the GFP. In the case of downregulation, GFP stability creates another issue that precludes detection of rapid effects. Altogether, this particular promoter library seems to be suboptimal for our experimental setup.

## Conclusions

In this work, we describe the transcriptional changes in *E. coli* during moderate and severe copper stress by RNA-seq. We detected the upregulation of known copper response pathways, histidine biosynthesis, oxidative stress response and iron acquisition pathways. During moderate copper stress, the bacteria retain better capacity to mount protective responses as we saw additional defensive pathways: biosynthesis of additional amino acids methionine and arginine, heat response and protein folding, lipid A biosynthesis activated compared to severe Cu stress. Additionally, we explored the use of a promoter-GFP library from Horizon Discovery to screen for transcriptional responses to copper. However, at least in our experimental setup, poor signal-to-noise ration made its use impractical and therefore we consider the RNA-seq results as a detailed view of copper stress response in *E. coli*.

## Supporting information

Supplementary file 1 - Enrichment analysis

Supplementary file 2 - RNAseq differentially expressed genes

## Acknowledgements

This work was supported by the Estonian Research Council grants PRG1496, TemTA55 and TK210 (AI), and STP39 (AA), and Horizon Europe project 101159721 (AI). This research is conducted by partially using the research infrastructure “Experimental Studies and Applications of Cellular Processes – RAKERA” funded by the Estonian Research Council (TARISTU24-TK14).

We thank Merilin Rosenberg for her valuable ideas and support.

## Author contributions

HA – conceptualization, study design, investigation, data analysis, interpretation and management, writing the manuscript; KJ – investigation, formal data analysis; AA – data analysis and interpretation, review and editing of the manuscript, funding acquisition; AI – conceptualization, study design, review and editing of the manuscript, project administration, funding acquisition

