## Supplementary Figures for "Copper stress upregulates oxidative stress response, histidine production and iron acquisition genes in *E. coli*"

### **Supplementary data**

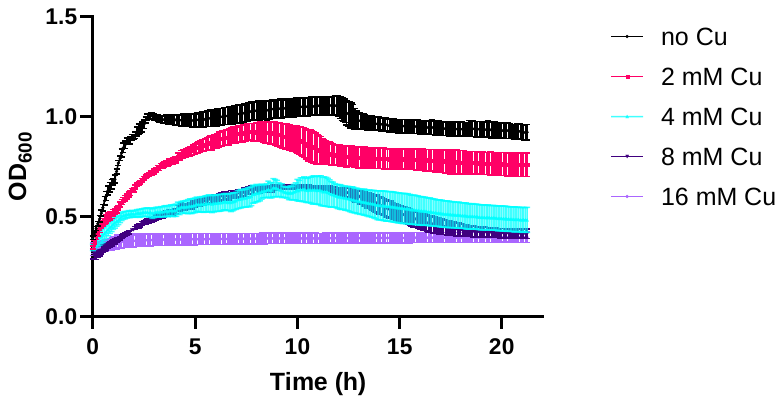

**Supplementary Figure 1.** Determining *E. coli* MG1655 MIC for the growth of transcriptome samples. The error bars represent mean ± SD of one experiment with four replicates.

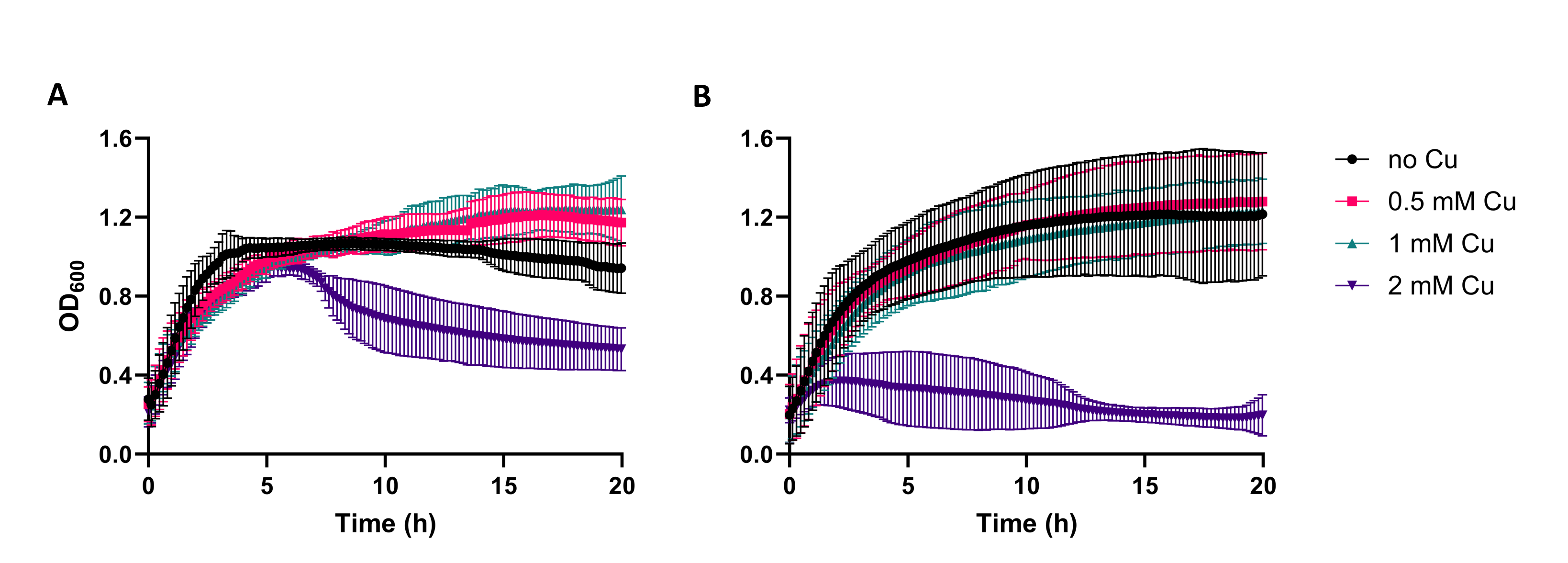

**Supplementary Figure 2.** Determination of the MIC of *E. coli* MG1655 carrying the pUA66‑fes plasmid (Km⁺) in MOPS (**A**) and LB (**B**). Error bars represent mean ± 95% CI of four biological experiments.
